# Exploring the potential of *Bacillus subtilis* as cell factory for food ingredients and special chemicals

**DOI:** 10.1101/2023.05.08.539805

**Authors:** Taichi Chen, Stanley Brul, Jeroen Hugenholtz

**Affiliations:** Molecular Biology and Microbial Food Safety, Swammerdam Institute for Life Sciences, University of Amsterdam, 1098 XH Amsterdam, The Netherlands; NoPalm Ingredients BV, Nieuwe Kanaal 7a, 6709 PA Wageningen, The Netherlands

**Keywords:** *Bacillus subtilis*, Fermentation, Primary metabolites

## Abstract

**Background:** *Bacillus subtilis* has been established as model microorganism for fundamental research in the laboratory on protein production/secretion and sporulation and as model bacterium for controlling spoilage in the food industry. It has also been used for production of (commercial) enzymes and several secondary metabolites such as vitamins. However, this doesn’t fully reflect the potential of *B. subtilis* as a cell-factory. Here, various strains of *B. subtilis*, including food-grade, spore-deficient strains and industrially used strains, were compared for their growth and metabolic potential. Industry-relevant parameters were analyzed for all strains under various aeration regimes, under anaerobic conditions, in various nutritious and nutrient-limited cultivation media, with and without organic nitrogen sources, and with and without sugar.

**Results:** Practical experiments were conducted to compare industrial relevant properties like growth rates, intracellular components and extracellular metabolite profile of different *B. subtilis* strains. Based on growth flexibility in different media, we found that some strains like NCIB3610 and DSM1092 are adapted to inorganic or organic nitrogen source utilization, which is highly relevant when considering a biorefinery approach using various cheap and abundant waste/sidestreams. Secondly, spore-deficient strains such as 3NA, 168S and PY79S, showed advantages in microbial protein and acetolactate pathway expression, which is associated with applications in food industry for protein supplement and diacetyl production. Lastly, WB800 and PY79S exhibited potential for fermentative production of Dipicolinic acid, 2,3-Butanediol and Lactic acid that could serve as precursors for biopolymers.

**Conclusion:** This study demonstrates the broad potential for more extensive industrial use of *Bacillus subtilis* in the (bio-based) chemical industry for use of sidestreams, in the personal care industry, in the food industry for food additive production, and in the bio-sustainable industry for biofuel and bio-degradable plastic precursors production. In addition, selecting different *B. subtilis* strains for specific purposes makes full use of the diversity of this species and increases the potential of *B. subtilis* in its contribution to the bio-based economy.

## Introduction

As the market for sustainable chemical production expands, researchers have been exploring bioproduction of chemicals for several decades. This includes the use of microbial fermentation to produce chemicals of interest. Compared with conventional chemical approach, microbial cell factories require lower temperature and pressure, and can convert renewable substrates into products of interest. Some model microorganisms like *Escherichia coli*[1]*, Bacillus subtilis*[2] and *Saccharomyces cerevisiae*[3] are promising candidates for industrial-relevant chemicals production due to their developed databases and extensive molecular toolbox.

*B. subtilis* is a representative gram-positive bacterium that has been intensively used in studies of sporulation process and protein production/excretion because of its robustness as a spore former and its excellent protein secretory capability[4]. In addition, it is recognized as a plant growth-promoting bacterium (PGPB), that enhances plant growth and protects plants from phytopathogen[5], instead of chemical fertilizers and pesticides, can contribute to an eco-friendly and sustainable economy. Moreover, different *B. subtilis* strains have been developed through mutagenesis or metabolic engineering to metabolize various carbon/nitrogen sources into several secondary metabolites such as (commercial) enzymes[6] and vitamins[7] or into biochemicals, such as oligosaccharides[8] and organic acids[9].

Among all *B. subtilis* strains, *B. subtilis* 168[10] is the major workhorse for versatile cell factory construction because of its sufficient gene annotations and high transformation efficiency. However, focusing solely on *B. subtilis* 168 could ignore the full potential of this species as many variants have unique advantages in environment adaption or metabolism (Fig. 1). For example, *B. subtilis* 3NA[11] can reach high cell density during fed-batch fermentation, which is an attractive point, but only limited work has been done to explore its potential. Also, *B. subtilis* PY79[12] owns a large amount of-omics data that can help metabolic rewiring, but it is mainly used in studies related to spores and biofilm[13]. Therefore, we propose that current research don’t fully reflect the potential of *B. subtilis* as a cell-factory, and systematic evaluation of metabolic potential of different *B. subtilis* strains may help expand the application of this species in more aspects in the bio-based industry (The difference and details of all selected *B. subtilis* strains are discussed in the “Strains and media” section).

**Figure 1.**
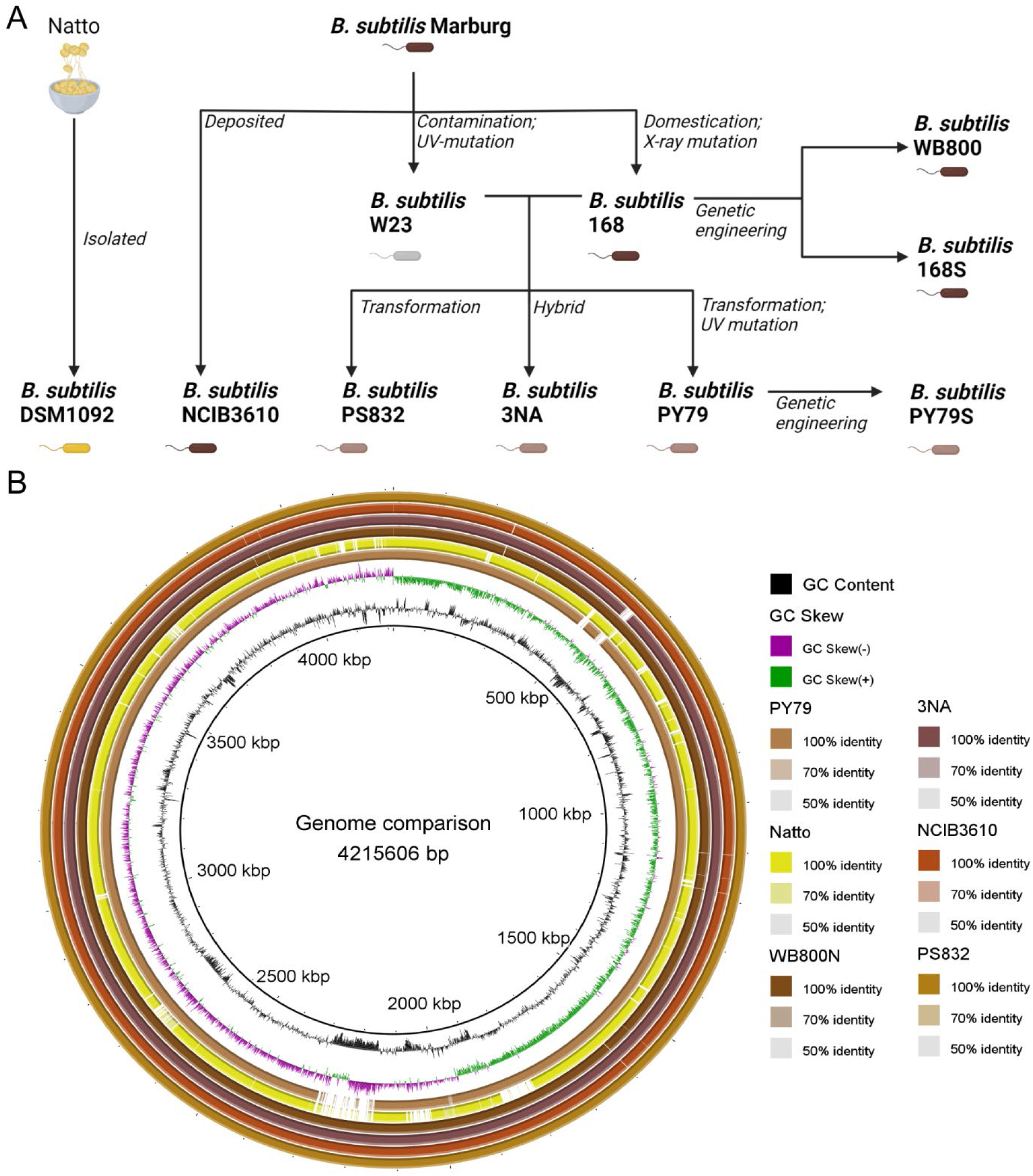
A. Genome heritage of nine *B. subtilis* strains. B. Genomic sequence comparison of representative *B. subtilis* strains using genome of 168 as the reference.

In this research, we investigated metabolic potential of nine *B. subtilis* variants in view of extending the use of this model species. First, we assessed the metabolic flexibility of *B. subtilis* strains regarding various media. Second, the effect of dissolved oxygen on growth and extracellular metabolites accumulation were investigated in flask level. Finally, we discussed the possibilities of different *B. subtilis* strains acting as cell factories and make practical recommendations for strain selection for industrial applications for different purposes.

## Results

Variation in growth characteristics of *B. subtilis* strains associated with medium To investigate the metabolic flexibility on different nitrogen sources of *B. subtilis* strains, we cultivated nine *B. subtilis* strains on different media in 96-well plates and plotted their specific growth rates (µ) and growth curves in Fig. 2. The specific growth rates of the strains were pairwise compared by one-way ANOVA (Supplementary Fig 1).

**Figure 2.**
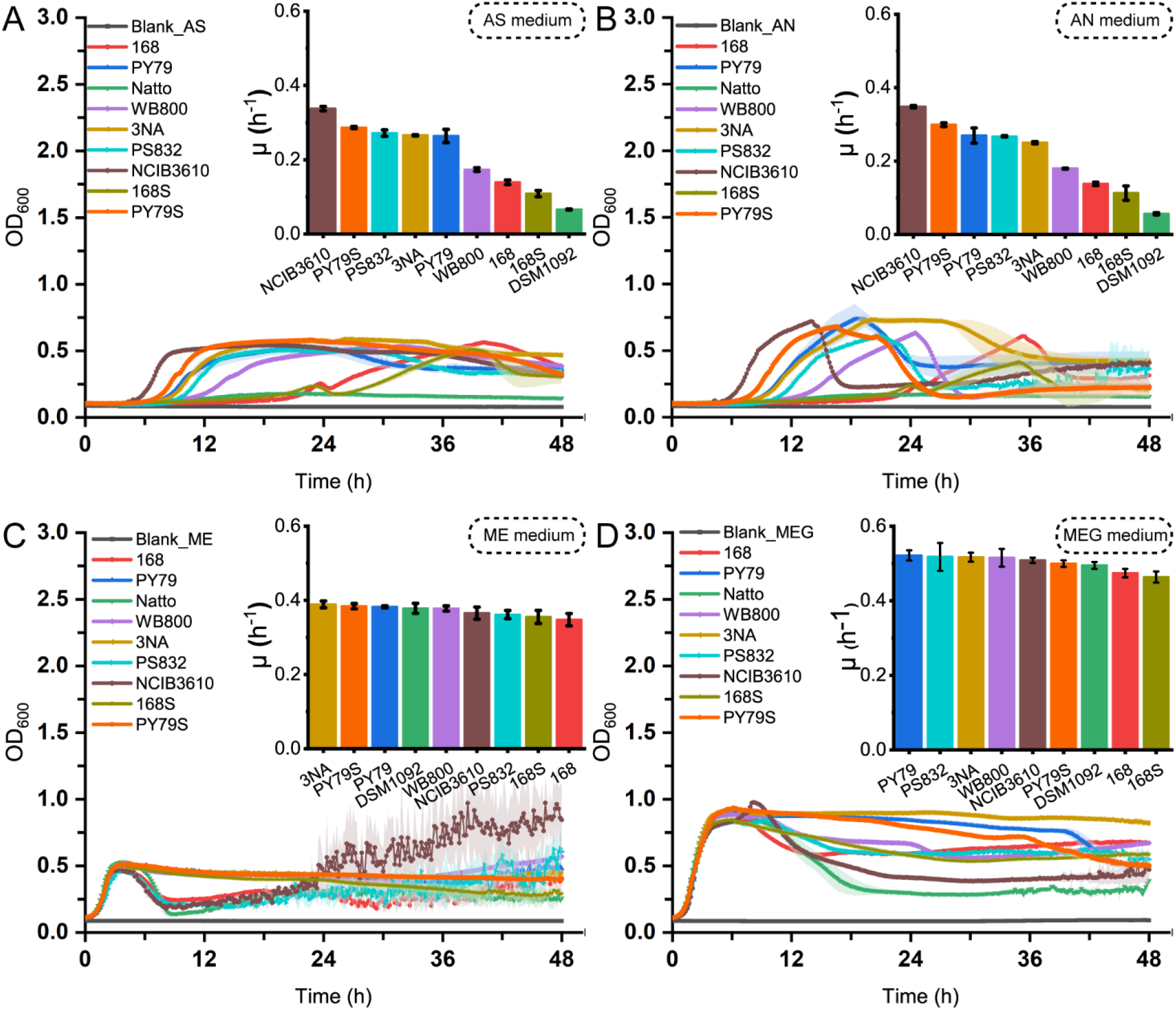
Growth curves and specific growth rates of nine *B. subtilis* strains on various media. A. Growth properties of nine strains in medium with ammonium sulphate; B. Growth properties of nine strains in medium with ammonium nitrate; C. Growth properties of nine strains in medium with meat extract; D. Growth properties of nine strains in medium with meat extract and glucose. The shallow surrounding growth curve represents the standard deviations of replicates.

*B. subtilis* NCIB3610[14], PY79S, PS832[15], 3NA and PY79 exhibited higher specific growth rates than other strains in media with ammonium salts (Fig. 2A, Fig. 2B), suggesting that they are competitive in these environments. PS832, 3NA and PY79 showed no significant difference in specific growth rate in the media containing ammonium salts. *B. subtilis* NCIB3610, however, showed significant difference with other strains with the highest growth rate of 0.33 h^-1^ and 0.35 h^-1^ among all strains in media with ammonium sulfate and with ammonium nitrate, respectively (Fig. 2A, Fig. 2B). Subsequently, we replaced the ammonium salts and glucose with meat extract (ME medium) to assess the growth characteristics of *B. subtilis* strains in the environment containing free amino acids, peptides and proteins. In ME medium, most strains showed no significant difference in specific growth rates with others. Notably, the growth rate of DSM1092[16] increased from 0.06 h^-1^ to 0.38 h^-1^ after switching the nitrogen source from inorganic to organic, suggesting that it is adapted for the environment containing organic nitrogen source (Fig. 2C). By further adding glucose into ME medium (MEG medium), we observed that the growth rate of DSM1092 increased by 28.9% from 0.38 h^-1^ to 0.49 h^-1^ (Fig. 2D).

In addition to growth rates, *B. subtilis* strains exhibited different growth patterns when growing in different media. In the media containing organic nitrogen sources, most strains showed immediate propagation and started to decline after reaching the peak, whereas the downward trend was not the same for different strains (Fig. 2C, Fig. 2D). In media with ammonium salts, all strains presented obvious lag phases (Supplementary table 1) and long stationary phase when growing in the medium with ammonium sulfate (AS medium) (Fig. 2A). Unlike the abovementioned cases, strains showed both different lag phases and a decreasing trend after reaching the highest biomass in the medium with ammonium nitrate (AN medium) (Fig. 2B).

### Variation in growth pattern of *B. subtilis* strains associated with oxygen

To evaluate the effect of oxygen on *B. subtilis* growth, we created high Dissolved oxygen (DO), optimal DO and oxygen-deprived conditions for fermentation experiments in flask and measured OD_600_ values of strains in 72h. When ammonium salt was used as the only nitrogen source, only the growth of NCIB3610 showed an obvious improvement with increase of dissolved oxygen(Fig. 3A, Fig. 3B). However, in the medium containing free amino acids, all strains grew faster with the increase of dissolved oxygen. For example, in the medium with meat extract and glucose (MEG medium), the OD_600_ value of *B. subtilis* DSM1092 reached 16, which was 3 times the value under optimal DO condition (Fig. 3D). Furthermore, as the protein and polypeptide content in the medium decreased, the influence of dissolved oxygen on the growth pattern of the strain became less. When casein hydrolysate was the only organic nutrient in the medium, elevating dissolved oxygen had no effect on the growth of all strains (Supplementary Fig. 2C).

**Figure 3.**
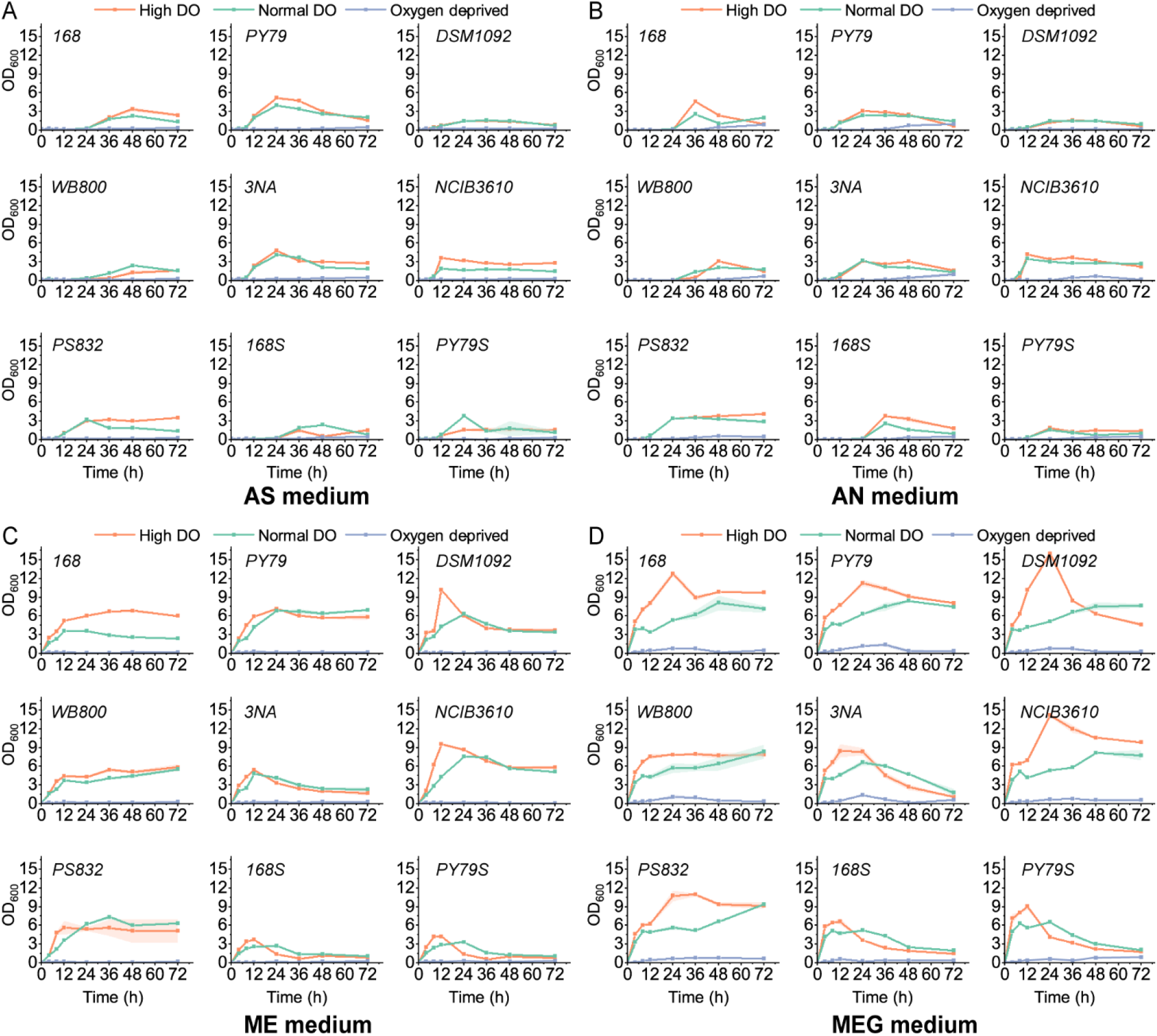
A∼D. Growth curves of strains in flasks in medium with ammonium sulphate (AS), ammonium nitrate (AN), meat extract (ME) and both meat extract and glucose (MEG) under high DO, optimal DO and oxygen-deprived conditions. The shallow surrounding growth curve represents the standard deviations of replicates.

Additionally, we observed that *B. subtilis* strains only grew in the presence of both glucose and nitrate or both glucose and organic nitrogen sources under oxygen-deprived condition. However, NCIB3610 reached 0.5 in the medium with ammonium nitrate (AN medium), far exceeding other variants (Fig. 3B). Moreover, 3NA and PY79 whose highest OD_600_ were both 1.4, outpaced other strains by faster growth and higher biomass in MEG medium (Fig. 3D).

### Variation in intracellular components of *B. subtilis* strains

After comparing growth flexibility of strains under different conditions, we turned to cellular components of different strains, such as microbial protein and spores. To compare the protein in biomass and dry cell weight of *B. subtilis* strains, we conducted fermentation experiments in LB medium supplied with glucose (LBG medium) in 250-ml flasks for 24h. At 6h (exponential phase), *B. subtilis* strains showed similar dry cell weight and protein proportions, while the protein ratios of strains varied from 11% to 38% at 24h (stationary phase) (Table 1). At 24h, spore-deficient *B. subtilis* 3NA exhibited the highest proportion of 38%, 3.4 times higher than that of PY79 which is a well-known spore-forming variant. The protein proportion of PY79 also increased from 11% to 36% after we disrupted its sporulation via knocking out *spo0A* (PY79S), indicating the negative effect of sporulation on microbial protein proportion.

**Table 1.** Dry cell weight and protein concentration of strains per OD_600_.

| Strains | 6h |  |  | 24h |  |  |
| --- | --- | --- | --- | --- | --- | --- |
|  | Dry weight (mg/l) | Protein concentration (mg/l) | Proportion Protein/Dry weight (%) | Dry weight (mg/l) | Protein concentration (mg/l) | Proportion Protein/Dry weight (%) |
| MG1655 | 530 ± 23 | 89 ± 9 | 17 ± 1 | 288 ± 4 | 113 ± 15 | 39 ± 6 |
| 168 | 415 ± 30 | 48 ± 14 | 11 ± 3 | 425 ± 42 | 77 ± 10 | 18 ± 1 |
| PY79 | 390 ± 41 | 57 ± 9 | 14 ± 2 | 500 ± 16 | 53 ± 6 | 11 ± 1 |
| DSM1092 | 355 ± 122 | 68 ± 12 | 21 ± 7 | 538 ± 31 | 82 ± 5 | 15 ± 0 |
| WB800 | 400 ± 67 | 58 ± 17 | 21 ± 10 | 400 ± 8 | 123 ± 24 | 31 ± 8 |
| 3NA | 415 ± 33 | 68 ± 5 | 16 ± 0 | 325 ± 47 | 122 ± 16 | 38 ± 12 |
| NCIB3610 | 440 ± 12 | 66 ± 8 | 15 ± 2 | 260 ± 69 | 66 ± 9 | 27 ± 6 |
| PS832 | 435 ± 18 | 73 ± 5 | 17 ± 1 | 210 ± 56 | 38 ± 5 | 19 ± 3 |
| 168S | 460 ± 34 | 72 ± 9 | 16 ± 1 | 588 ± 37 | 166 ± 2 | 28 ± 2 |
| PY79S | 420 ± 32 | 51 ± 12 | 12 ± 3 | 475 ± 15 | 171 ± 2 | 36 ± 1 |

Although *B. subtilis* spores are usually discussed with food spoilage and contamination, they are also related to dipicolinic acid (DPA) production which is industrial relevant. To determine spore dry weight and DPA content in the spores of *B. subtilis* strains, we cultivated strains in MOPS medium in 250-ml flasks for 72h, and purified spores. As a result, we didn’t find DPA in supernatant of cultures but only in spores. *B. subtilis* 168 stored 74 mg of DPA in 1 g of dry spore, which is the lowest among selected strains (Table 2). However, two mutants WB800[17] and PS832 derived from 168 showed the highest DPA production, namely 128 mg and 126 mg of DPA in 1 g of dry spore, respectively.

**Table 2.** Spore properties of strains.

| Strains | Dry weight (g/l/OD) | Dipicolinic acid concentration (mg/l/OD) | Proportion (mg<sub>DPA</sub>/g<sub>spore</sub>) |
| --- | --- | --- | --- |
| 168 | 0.09 ± 0.00 | 6.68 ± 0.19 | 74.56 ± 0.87 |
| PY79 | 0.11 ± 0.01 | 10.22 ± 0.16 | 91.13 ± 8.25 |
| DSM1092 | 0.13 ± 0.00 | 7.56 ± 0.15 | 56.36 ± 2.43 |
| WB800 | 0.10 ± 0.01 | 12.29 ± 0.21 | 127.59 ± 10.32 |
| NCIB3610 | 0.10 ± 0.00 | 8.77 ± 0.12 | 90.77 ± 0.66 |
| PS832 | 0.05 ± 0.00 | 6.55 ± 0.20 | 126.34 ± 12.88 |

### Variation in extracellular metabolites distribution of *B. subtilis* strains

To further investigate the metabolic distribution during fermentation, the profiles of organic acids and some primary metabolites were determined. Firstly, we observed pathway preference of *B. subtilis* strains over time. For example, under optimal Dissolved Oxygen (DO) condition, the acetolactate pathway had a greater flux than other two pathways at 12h in DSM1092 in MEG medium while the acetate synthesis pathway took over at 24h (Fig. 4A). Under high DO condition, however, lactate production outcompeted other two directions at 12h and we didn’t see intermediate in the acetolactate pathway at 24h.

**Figure 4.**
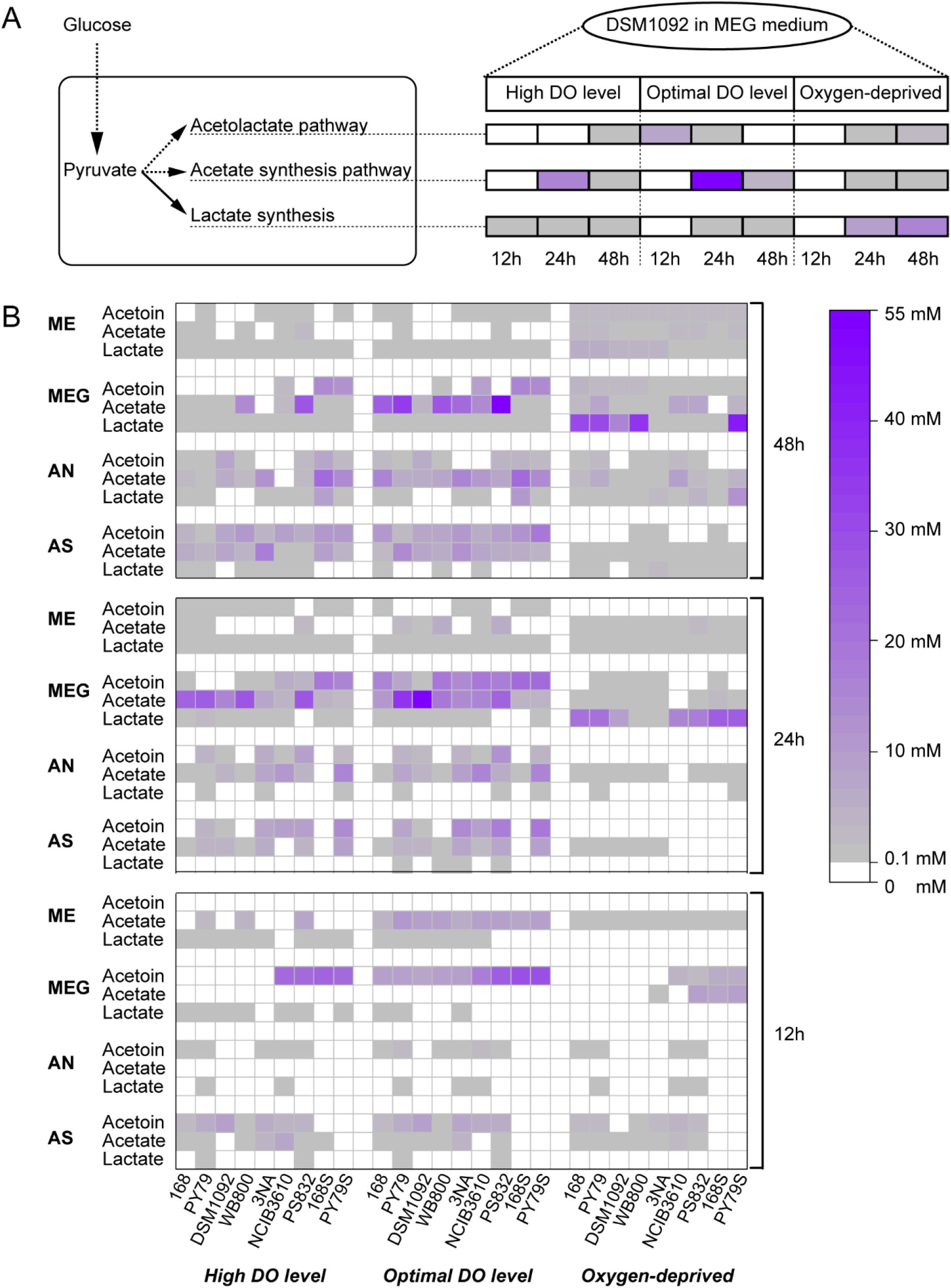
A. Metabolite profile of DSM1092 in MEG medium. B: Metabolic distribution of nine *B. subtilis* strains in medium with ammonium sulphate (AS), ammonium nitrate (AN), meat extract (ME) and both meat extract and glucose (MEG) under high DO, optimal DO and oxygen-deprived conditions.

We take acetoin as an indicator for the acetolactate pathway because acetoin is the main intermediate in this pathway, which also leads to Diacetyl and 2,3-Butanediol production. Under aerobic conditions (Fig. 4B), acetoin was mainly produced in the first 24h and was converted later (to butanediol) in ME and MEG medium but slowly accumulated in AS medium under aerobic conditions. An interesting finding is that *B. subtilis* strains produced the lowest titers of acetoin in ME medium in which glucose was absent, suggesting an important role for glucose in the acetolactate pathway. 168S and PY79S producing the most acetoin both at 29 mM in MEG medium while 168, PY79, DSM1092, 3NA and PS832 depleted all acetoin after 48h in MEG medium (Fig. 4B).

Then we assessed the acetate synthesis pathway. The most interesting finding was two different patterns of acetate production between *B. subtilis* DSM1092 and PS832 in MEG medium under aerobic conditions. Acetate concentrations in the supernatants of PS832 fermentation were 0, 24.1 and 49.7 mM in 3 days, respectively (Fig. 4B). In contrast, the corresponding data in the supernatant of DSM1092 fermentation were 0, 48.4 and 4.1 mM, respectively. The continuous accumulation of acetate by PS832 indicated the risk of wasting preferred carbon sources to by-products during fermentation while DSM1092 showed the potential for growth on overflow metabolites.

In addition to the above two pathways that were strongly expressed under aerobic conditions, we also observed that under oxygen-deprived condition, lactate is the main product of most strains due to NAD^+^ regeneration during lactate synthesis. In all conditions, *B. subtilis* PY79S produced 42.36 mM lactate in MEG medium which was the highest among variants (Fig. 4B).

## Discussion

In this study, we investigated metabolic flexibility of different *B. subtilis* strains corresponding to various industrial applications and recommended certain variants for specific usage (Table 3).

**Table 3.**
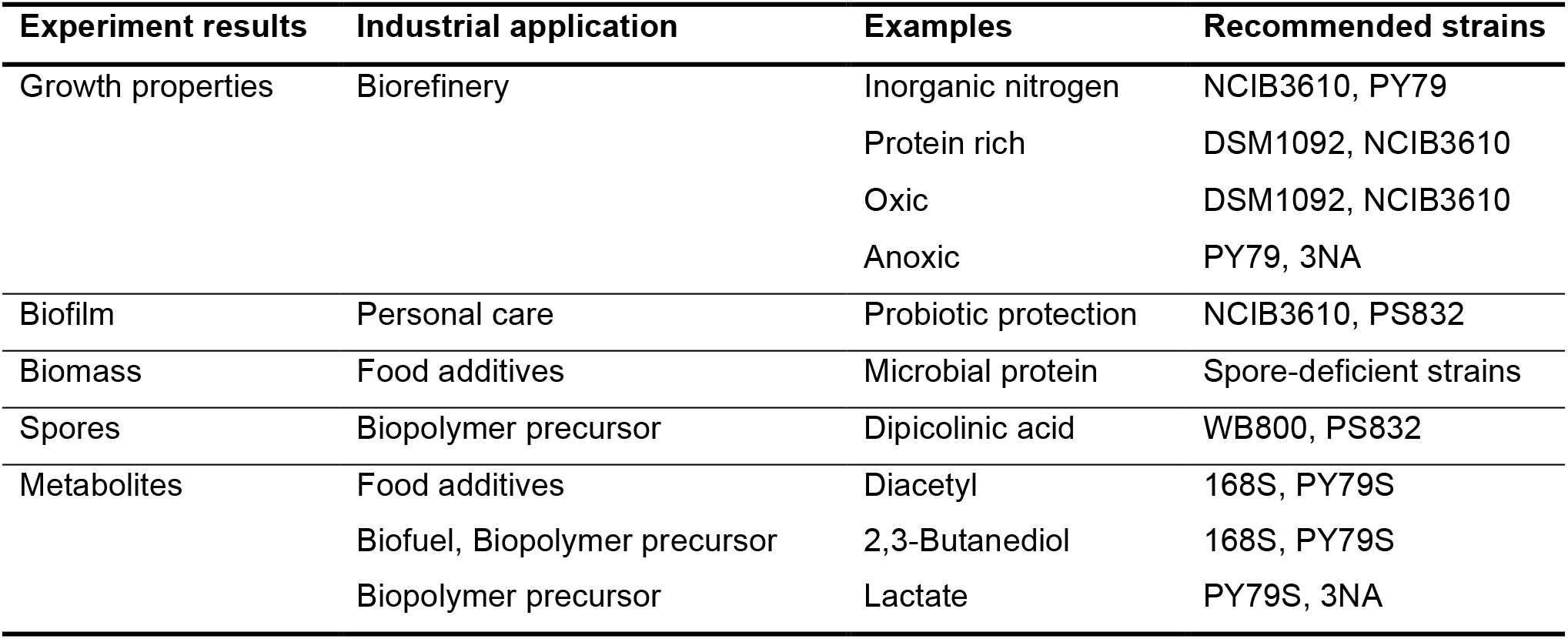
Proposed industrial applications based on experiments.

### Biorefinery

For cost-effective production of bio-based chemicals and (food) ingredients, it is essential that cell-factories can be productive based on cheap and abundantly available biomass, waste- and sidestreams. Using such feedstocks, the preferred cell-factories will need to cope with large variations in nutrient composition, with respect to sugar content, nitrogen content, in oxygen supply and in other parameters such as low or high pH, presence of anti-nutritional factors, etc.

What’s more, we also observed various duration of lag phase of different strains (Supplementary Table 1). Although the lag phase can be avoided by seed cultivation in industry, the shortest lag phase duration of *B. subtilis* NCIB3610 in both AS and AN medium suggested that it’s more adapted to the nutrient-limited environments than other strains. Given those advantages with the high growth rate, *B. subtilis* NCIB3610 seemed to be the most promising host for aquaculture wastewater denitrification.

On the other hand, DSM1092 and some *B. subtilis* 168 legacy strains (Fig. 1A), like WB800, and 168S showed difficulty in growing on media with sole inorganic nitrogen source. But their growth was obviously improved by addition of protein and amino acids especially when DO level was increased (Fig. 3C, Fig. 3D).

However, DSM1092 in this study exhibited rapid depletion of overflow compounds (acetate and acetoin) in MEG medium without an intact glyoxylate shunt reported (Fig. 4A). This characteristic is beneficial for industrial applications, such as growth on additional carbon sources and better robustness towards excess glucose in environment.

Based on the growth properties of strains, we hypothesize that a) Strains containing part of genome sequence of *B. subtilis* W23 are more adapted to different nutritional environments (Fig. 1B). b) *B. subtilis* DSM1092 and some legacy strains of 168 have domesticated metabolic pathways for organic nitrogen sources utilization rather than ammonium salts because there are always sufficient organic nitrogen sources in their culture environment.

### Pellicle formation

The biofilm matrix of *B. subtilis* has been extensively studied for various personal care applications such as biocatalyst for therapeutic Menaquinone-7 production or as protector for beneficial probiotics[18][19]. Among all selected variants, *B. subtilis* NCIB3610 and PS832 formed clear pellicles in media containing organic nitrogen source at first day (Supplementary Fig. 1), suggesting that *B. subtilis* NCIB3610 and PS832 are more suitable for biofilm matrix producing than others. On the other hand, biofilm can be a disadvantage during liquid state fermentation, as it affects the transfer of nutrients and oxygen in the system and increase the difficulty of fermenter cleaning. In this regard, *B. subtilis* 168S is a better choice to avoid the biofilm matrix during fermentation as its biofilm formation capacity may be disturbed by deletion of *spo0A*.

### Food additives

Current protein production for feed and food, involving various plant and animals, is highly inefficient, has major environmental impact with respect to land/water-use and Greenhouse gas emissions and often involves non-local production resulting in a need for long-distance transportation. However, the microbes offer a more efficient approach to obtaining protein as they are able to grow fast (hours to double their biomass), have innately high protein content and can be produced locally with limited land- and water-usage. In our case, all spore-deficient variants (*B. subtilis* 3NA, 168S and PY79S) showed higher protein proportion than others (∼37% by dry cell weight) while the famous spore producer - PY79 showed the lowest protein ratio. These findings suggested that the spore-deficient *B. subtilis* strains can serve as good microbial protein producers.

Diacetyl is a flavour responsible for buttery aroma and is widely used in the food and beverage industry[20]. It can be transformed spontaneously from acetoin in the acetolactate pathway, so extracellular acetoin profiles can reflect the capacity of Diacetyl production of *B. subtilis* variants. Based on metabolite profiles, we observed that optimal DO conditions were conducive to the expression of acetolactate pathway while high dissolved DO promoted biomass gaining. Given the growth and metabolites profile in AS medium, we proposed that the non-sporulate mutants (especially PY79S) might be good hosts for potential products within the acetolactate pathway such as diacetyl because they had shorter fermentation time, hence, higher productivity.

### Bio-based polymers

Over past decades, the increasing demand for “green materials” have opened new windows of opportunity for bio-based products like bio-based polymers that is more environmental-friendly than petroleum-derived consumables.

Dipicolinic acid, Lactic acid and 2,3-Butanediol are promising monomers which can be harnessed by fermentation and can be used for polymeric materials production. Researchers have been developing different microorganism to covert sugars into DPA. For example, Lynn et al. established a DPA synthesis pathway by overexpression of dipicolinate synthase genes from *Paenibacillus sonchi* in a *Corynebacterium glutamicum* L-lysine producer strain, resulting in 2.5 g/l of DPA in shake-flask[21]. K. McClintock et al. achieved a titer of 4.7 g/l DPA using an *E. coli* strain by knocking out competitive pathways and heterogeneous expressing dipicolinate synthase from *B. subtilis*[22]. Compared with abovementioned bacteria, *B. subtilis* possesses complete DPA synthesis pathway naturally as DPA is a vital compound in *B. subtilis* spores for heat resistance. In addition, *B. subtilis* can also produce L-lactic acid which is a bulk chemical and can be polymerized to produce biodegradable polymer PLA for food packaging[23]. Combining the growth and metabolite profiles of *B. subtilis* strains under different DO conditions, we recommended WB800 and PS832 as hosts for DPA production under aerobic conditions, PY79S as promising host for 2.3-Butanediol and lactate production under aerobic conditions and oxygen-deprived condition, respectively.

## Conclusion

In this study, we first investigated the growth flexibility of various *B. subtilis* strains under different conditions, such as growth rates and cellular components. Then, we elucidated the effects of environment on the metabolism of *B. subtilis* strains, such as the distribution of metabolites in the acetolactate pathway, lactate synthesis pathway and acetate synthesis pathway in different periods. Based on results, we recommend some *B. subtilis* strains that are uncommonly used in the development of cell factories, such as DSM1092 and 3NA for further development of cell factories. All in all, we conclude that *B. subtilis* is an underdeveloped cell factory with great metabolic flexibility under different nutrient conditions and various aeration regimes. By understanding its metabolic potential under various conditions, we can expand the use of *B. subtilis* beyond enzyme production.

## Material and methods

### Strains and media

All strains used in this study are listed in Table 4. As shown in Fig. 1A, *B. subtilis* NCIB3610 is the closest descendant of the lab strain *B. subtilis* Marburg and shares a similar genotype with the widely used transformation-efficient *B. subtilis* 168 (Fig. 1B). However, they have obvious differences in phenotypes like substrate utilization, metabolites synthesis and swarm ability due to the mutations at *trpC*, *swrA*, *sfp* and *gudB* loci. *B. subtilis* WB800, a derivative of *B. subtilis* 168, lacks eight extracellular proteases that helps to stabilize secreted proteins. *B. subtilis* PY79, 3NA and PS832 possess some genomic characteristics of *B. subtilis* W23 and have been commonly used for sporulation and germination research. The prototroph *B. subtilis* PY79 possesses 4.3% reduction in genomic size compared with *B. subtilis* 168, and 34 kb replacement of *B. subtilis* 168 genes with orthologous *B. subtilis* W23 genes. *B. subtilis* 3NA produces no spores due to a frame-shift mutation at *spo0A* locus and an elongation at *abrB* gene which both affects the initialization of sporulation. The *B. subtilis* PS832 is seemed as the “prototrophic and revertant” version of *B. subtilis* 168. It has been used to exploit spore germination like germinant receptors in Setlow’s lab and our lab for a long period. To address the potential demand for blocking sporulation, we also constructed two spore-deficient strains named *B. subtilis* 168S and *B. subtilis* PY79S in this study, whose *spo0A* gene was entirely knocked out. In addition to the family of lab strains, we selected one food-grade *B. subtilis* DSM1092 that is isolated from traditional Japanese food Natto for research .Overall, we selected nine representative *B. subtilis* strains to explore their metabolic potential for cell factory development.

**Table 4.** Strains used in this study.

| Strains | Description (; source) | Reference |
| --- | --- | --- |
| *B. subtilis* NCIB3610 | Wild type; BGSC<sup>a</sup> | [14] |
| *B. subtilis* 168 | *trpC2*; Lab stock | [10] |
| *B. subtilis* PY79 | Prototroph SPβ<sup>s</sup>; Lab stock | [12] |
| *B. subtilis* DSM1092 | Wild type; DSMZ<sup>b</sup> | [16] |
| *B. subtilis* WB800 | *trpC2, nprE, aprE, epr, bpr, mpr::ble nprB::bsr Δvpr wprA::hyg*;<br>Lab stock | [17] |
| *B. subtilis* 3NA | *spo0A3*; BGSC | [11] |
| *B. subtilis* PS832 | Prototroph; Lab stock | [15] |
| *B. subtilis* 168S | *B. subtilis* 168 $\Delta spo0A :: Cm^R$; Lab stock | This study |
| *B. subtilis* PY79S | *B. subtilis* PY79 $\Delta spo0A :: Cm^R$; Lab stock | This study |
| *E. coli* MG1655 | F<sup>-</sup>, λ<sup>-</sup>, *rph-1*; Lab stock | [24] |

<sup>a</sup>BGSC Bacillus genetic stock center; <sup>b</sup>DSMZ Leibniz institute DSMZ-German collection of microorganisms and cell cultures GmbH

The Luria-Bertani (LB) medium was used for strain propagation and consisted of 5 g/l Yeast extract, 10 g/l Tryptone, and 5 g/l NaCl. For seed culture and microbial protein fermentation, LBG was adapted with the addition of 1‰ 50%(w/v) glucose in LB medium. All fermentation media contained the same basal salts solution: 12.5 g/l K_2_HPO_4_·3H_2_O, 3 g/l KH_2_PO_4_, 3 g/l MgSO_4_·7H_2_O, 10 ml/l Trace elements solution. Trace elements solution consists of 4 g/l FeSO_4_·7H_2_O, 4 g/l CaCl_2_, 1 g/l MnSO_4_·5H_2_O, 0.04 g/l CoCl_2_·6H_2_O, 0.2 g/l NaMoO_4_·2H_2_O, 0.2 g/l ZnSO_4_·7H_2_O, 0.1 g/l AlCl_3_·6H_2_O, 0.1 g/l CuCl_2_·H2O, 0.05 g/l H_3_BO_4_. For strain cultivation in Meat extract (ME) medium, 9 g/l Meat extract was supplemented as the only nitrogen source. Similarly, 9 g/l (NH_4_)_2_SO_4_, NH_4_NO_3_, NH_4_Cl, peptone and casein hydrolysate were added as nitrogen sources in Ammonium sulfate (AS) medium, Ammonium nitrate (AN) medium, Ammonium chloride (ACl) medium, Peptone (PE) medium and Casein hydrolysate (CA) medium, respectively. By supplementing ME, PE and CA with an additional 6 g/l glucose, we made MEG, PEG and CAG media, respectively. In addition, AS, AN and ACl medium were supplemented with 6 g/l glucose as the carbon source. Particularly, 0.05 g/l of tryptophan was added in the media for 168, WB800 and 168S as they are auxotroph mutants. To wash strains, a 10 × PBS solution was prepared with 35.81 g/l Na_2_HPO_4_.12H_2_O, 2.45 g/l KH_2_PO_4_, 80.07 g/l NaCl, 2.01 g/l KCl.

### Construction of spore-deficient strains

The primers and DNA templates used for gene amplification are listed in Supplementary table 2. The homogenous arms of *spo0A*-gene were amplified from *B. subtilis* 168 and the chloramphenicol resistance gene was amplified from plasmid pDG1662 according to the manufacturer’s instruction (Takara Primestar max DNA polymerase). These three fragments were combined by overlapping Polymerase Chain Reaction (PCR) and purified by DNA purification kit (Thermofisher). The purified product was then transformed into *B. subtilis* 168 and PY79 by the method of Anagnostopoulos & Spizizen, resulting in the 168S and PY79S strains[25].

### Seed culture for fermentation

Cultures of strains were taken from glycerol stocks stored at -80 °C and were streaked on LB medium plates which were then placed in a 37 °C incubator overnight. For each strain, one colony was picked into a 50 ml Erlenmeyer flask loaded with 12 ml of LB medium. After 6 hours of incubation, the cultures were centrifuged at 10,000 rpm for 2 minutes, and the resulting pellets were washed three times with 1 × PBS. Finally, the pellets were resuspended in 1 × PBS to obtain OD of 10.

### Cultivation experiments in 96-well plate and flask

For the fermentation experiments in 96-well plates, each well was loaded with 150 µl medium and inoculated with a washed seed culture (1% (v/v)). The inoculated microplates were covered with oxygen-permeable films and inserted into the microplate reader for 48 h growth curve measurement at 37°C.

For the fermentation experiments in flasks, different dissolved oxygen (DO) conditions were created by loading different amount of fermentation broth in Erlenmeyer flasks or glass bottles[26]: 50 ml/250 ml flask (High DO condition), 100 ml/250 ml flask (optimal DO condition), and 18 ml/20 ml bottle (oxygen-deprived condition). After inoculation, the flasks were incubated in a shaking incubator at 37°C, 200 rpm for 72 hours. The bottles were placed static in a 37°C incubator. Samples were taken at 4, 8, 12, 24, 36, 48 and 72 hours.

### Dry cell weight and protein concentration measurement

The LBG medium was selected for strain propagation and the measurement of biomass was performed as follows. A single colony was picked up from LB medium plate into a flask containing 50 ml liquid LB medium with 50 μl glucose (50% solution) and placed in the shaking incubator at 37°C. After 6 and 24 hours, the cells were harvested and centrifuged at 10000 rpm for 2 minutes. After three washes with PBS solution, the OD_600_ of suspended culture was measured. The culture was then diluted or condensed to create a 2 ml suspension in PBS solution or Ambic water (ammonium bicarbonate with 1% SDS) with an OD of 4 for dry cell weight and protein concentration, respectively.

A piece of filter paper was used to filter 2ml of cell suspension with an OD of 4 in PBS. After filtering all the liquid, the filter papers were placed back in the stove at 95°C. For dry cell weight measurement, 10 pieces of filter paper (0.22 µm) were thoroughly dried in a 95°C stove and their weight was determined up to four decimal places. A piece of filter paper was used to filter 2ml of the cell suspension of OD = 4 in PBS. After all the liquid was filtered, the filter papers were placed back in the stove at 95°C. After every 24 hours of drying, the papers were weighed until the weight showed no change. For microbial protein measurement, the cells were lysed with glass beans (0.1 mm) and the tubes were centrifuged at 15,000 rpm for 10 minutes. Protein concentration of *B. subtilis* cells was measured using a Thermo Scientific™ Pierce™ BCA protein assay. All steps followed the instruction on the assay kit manual.

### Dry spore weight and Dipicolinic acid measurement

All strains were revived on LB agar plates and incubated at 37°C overnight. Single colony was picked up into 50 ml 3-(N-morpholino) propane sulfonic acid medium (MOPS) buffered defined liquid medium (Sigma-Aldrich, St. Louis, MI, USA). For DSM1092, which could not propagate in the MOPS medium, 1% (v/v) LB medium was added to the mixture. Pure spores of strains were harvested after 3-day incubation in a shaking incubator at 37°C, 200 rpm. The MOPS medium and spore purification method was as described previously by Tu and co-workers [27]. Spore dry weights were determined by weighing overnight freeze-dried spores with an OD of 10. To measure DPA in each sample, 2 OD_600_ of spore was resuspended with a buffer consisting of 0.3 mM (NH_4_)_2_SO_4_, 6.6 mM KH_2_PO_4_,15 mM NaCl, 59.5 mM NaHCO_3_ and 35.2 mM Na_2_HPO_4_. Subsequently, the suspended spores were incubated at 100°C for 1 hour with a control set incubated at 37°C for 1 hour. After incubation, samples were centrifuged at 15,000 rpm for 2 minutes and 10μl of the supernatant was then added to 115μL of another buffer (1 mM Tris, 150 mM NaCl) with and without 0.8 mM terbium chloride (Sigma-Aldrich, St. Louis, MI, USA) for a 15-min incubation at 37°C. The fluorescence of the samples was measured by a Synergy Mx microplate reader (BioTek; 270-nm excitation; 545-nm reading; gain, 100) (Bad Friedrichshall, Germany). The background fluorescence (without terbium, incubated at 37°C) was subtracted from that of all the samples. A calibration curve of 0–140 mg/l DPA was used to calculate DPA concentrations of the sample.

### Extracellular metabolites measurement

The supernatant samples were centrifuged at 15,000 rpm for 10 min and filtered by 0.22 µm filters. The standards for the High-Performance Liquid Chromatography (HPLC) measurement of oxoglutarate, fumarate, malate, lactate, acetate, acetoin, glucose and ethanol were prepared with a set of concentration within 0-50 mM. All standards and samples were measured by HPLC using a Shimadzu instrument (LC-20AT, Prominence, Shimadzu) equipped with an Ion exclusion Rezex ROA-Organic Acid H+(8%) column (300 × 7.8 mm column, Phenomenex), along with a guard column (Phenomenex). Aqueous H_2_SO_4_ (5 mM) was used as mobile phase at a flow rate of 0.5 ml/min at 55 °C. A wavelength of 210 nm was used for calibration and analysis with the SPD-20A UV/VIS detector, and a cell temperature of 40°C was set for refractive index detector (RID 20A, Shimadzu).

### Statistical Analysis

All data are presented as the means ± standard deviation (SD). Three biological replicates were measured for biomass and spore weight measurement. Statistical analyses like Ordinary One-way ANOVA (Tukey) were carried out using OriginPro 2022 software and Prism GraphPad software.

## Abbreviations

AS medium: Fermentation medium with ammonium sulfate as sole nitrogen source
AN medium: Fermentation medium with ammonium nitrate as sole nitrogen source
ME medium: Fermentation medium with meat extract as sole nitrogen source
MEG medium: ME medium with extra glucose
DO: Dissolved oxygen
DPA: Dipicolinic acid
HPLC: High-Performance Liquid Chromatography

## Declarations

### Ethics approval and consent to participate

Not applicable.

### Consent for publication

Not applicable.

### Availability of data and materials

The datasets used and/or analyzed during the current study are available from the corresponding author on reasonable request.

### Authors’ contributions

Conceptualization, JH, SB and TC; performing experiments and analyzing data, TC; drafting original manuscript, TC; reviewing and editing, JH and SB. All authors read and approved the final manuscript.

## Acknowledgements

We acknowledge Dr. Meike Wortel and Pim van Leeuwen for the help for R scripts and metabolites detection through HPLC.

