## Supplementary file Tables and Figure for "Exploring the potential of *Bacillus subtilis* as cell factory for food ingredients and special chemicals"

**Supplementary Table 1** The highest $OD_{600}$ and lag phase of strains in media with ammonium salts

| Strains | AS medium | | AN medium | | ACl medium | |
|---|---|---|---|---|---|---|
| | Highest OD | Lag phase (h) | Highest OD | Lag phase (h) | Highest OD | Lag phase (h) |
| 168 | 0.48 ± 0.00 | 17.59 ± 0.44 | 0.53 ± 0.00 | 18.75 ± 0.14 | 0.50 ± 0.00 | 21.75 ± 0.30 |
| PY79 | 0.45 ± 0.04 | 6.21 ± 0.28 | 0.68 ± 0.08 | 7.18 ± 0.23 | 0.50 ± 0.01 | 7.24 ± 0.03 |
| DSM1092 | 0.10 ± 0.00 | 6.80 ± 0.37 | 0.10 ± 0.00 | 7.31 ± 0.05 | 0.10 ± 0.00 | 7.40 ± 0.23 |
| WB800 | 0.46 ± 0.00 | 8.47 ± 0.45 | 0.55 ± 0.00 | 10.52 ± 0.06 | 0.47 ± 0.00 | 11.29 ± 0.15 |
| 3NA | 0.51 ± 0.00 | 6.69 ± 0.06 | 0.66 ± 0.02 | 7.29 ± 0.19 | 0.52 ± 0.01 | 7.50 ± 0.18 |
| PS832 | 0.43 ± 0.00 | 7.86 ± 0.12 | 0.53 ± 0.00 | 9.02 ± 0.07 | 0.45 ± 0.01 | 8.83 ± 0.06 |
| NCIB3610 | 0.46 ± 0.00 | 3.86 ± 0.11 | 0.64 ± 0.01 | 4.35 ± 0.04 | 0.44 ± 0.00 | 9.22 ± 0.06 |
| 168S | 0.44 ± 0.00 | 21.45 ± 0.20 | 0.36 ± 0.06 | 38.06 ±17.63 | 0.45 ± 0.00 | 24.38 ± 0.80 |
| PY79S | 0.49 ± 0.00 | 5.63 ± 0.02 | 0.59 ± 0.00 | 5.91 ± 0.05 | 0.52 ± 0.00 | 6.49 ± 0.12 |

**Supplementary Table 2** Primers and DNA templates for gene amplification in this study

| Primers | Sequences (5'-3') | Template |
|---|---|---|
| 5pF_spo0AKO | GTGTTCTTGTCGTCGGATTTCATC | *B. subtilis*168 |
| 5pR_spo0AKO | TCAAACATGAGAATTCCACGTTTCTTCCTCCCCAAATGTAG | |
| 5pCm.F_spo0AKO | AGGAAGAAACGTGGAATTCTCATGTTTGACAGCTTATCATCGG | pDG1662[a] |
| 3pCm.R_spo0AKO | TCAGCGCAAACTTATAAAAGCCAGTCATTAGGCCTATCTGACA | |
| 3pF_spo0AKO | GACTGGCTTTTATAAGTTTGCGCTGATAAATAGGAGGC | *B. subtilis*168 |
| 3pR_spo0AKO | TCGAAATTCACGGTGCAAACGG | |
| spo0A.VF | GTTGAAAGAAACAGGAGGCATCGT | Mutants on |
| sop0A.VR | CCTAACTCTCCGTCGCTATTGTAACC | plate |

[a]This plasmid can be purchased in the Bacillus Genetic Stock Center

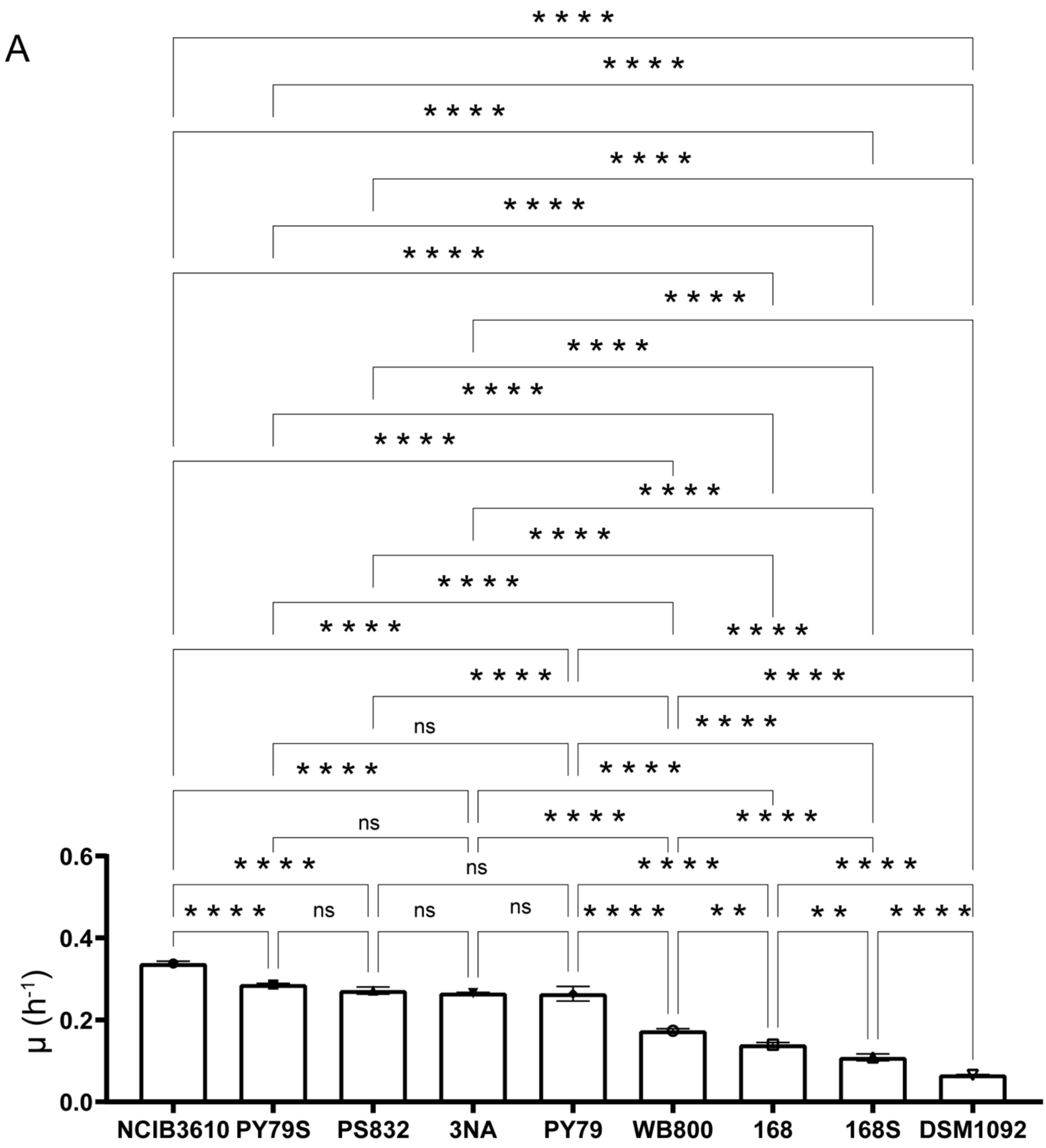
A
μ (h⁻¹)
0.0
0.2
0.4
0.6
NCIB3610
PY79S
PS832
3NA
PY79
WB800
168
168S
DSM1092
ns
* * * *
* *

B

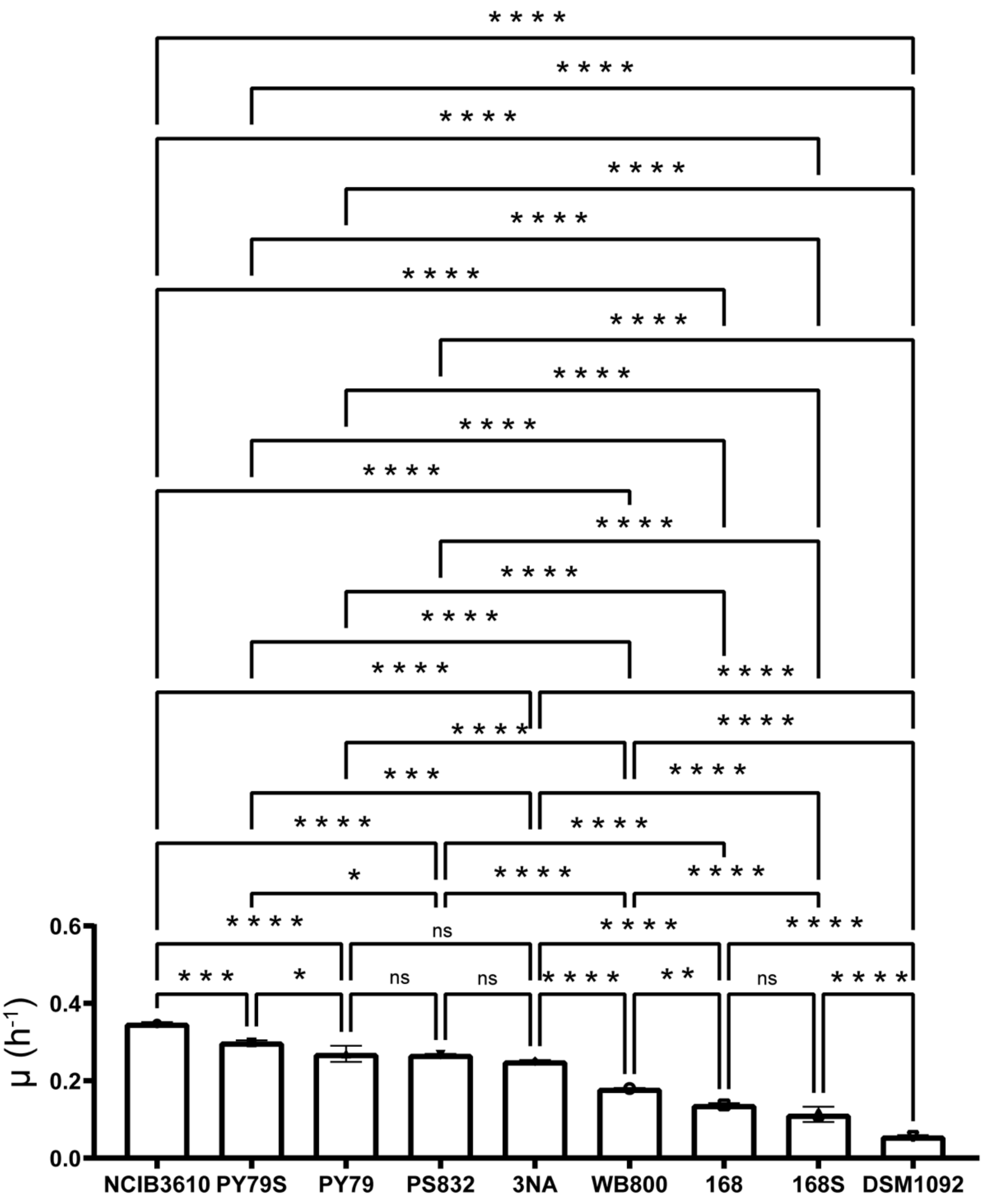

μ (h-1)
0.0
0.2
0.4
0.6
NCIB3610
PY79S
PY79
PS832
3NA
WB800
168
168S
DSM1092
ns

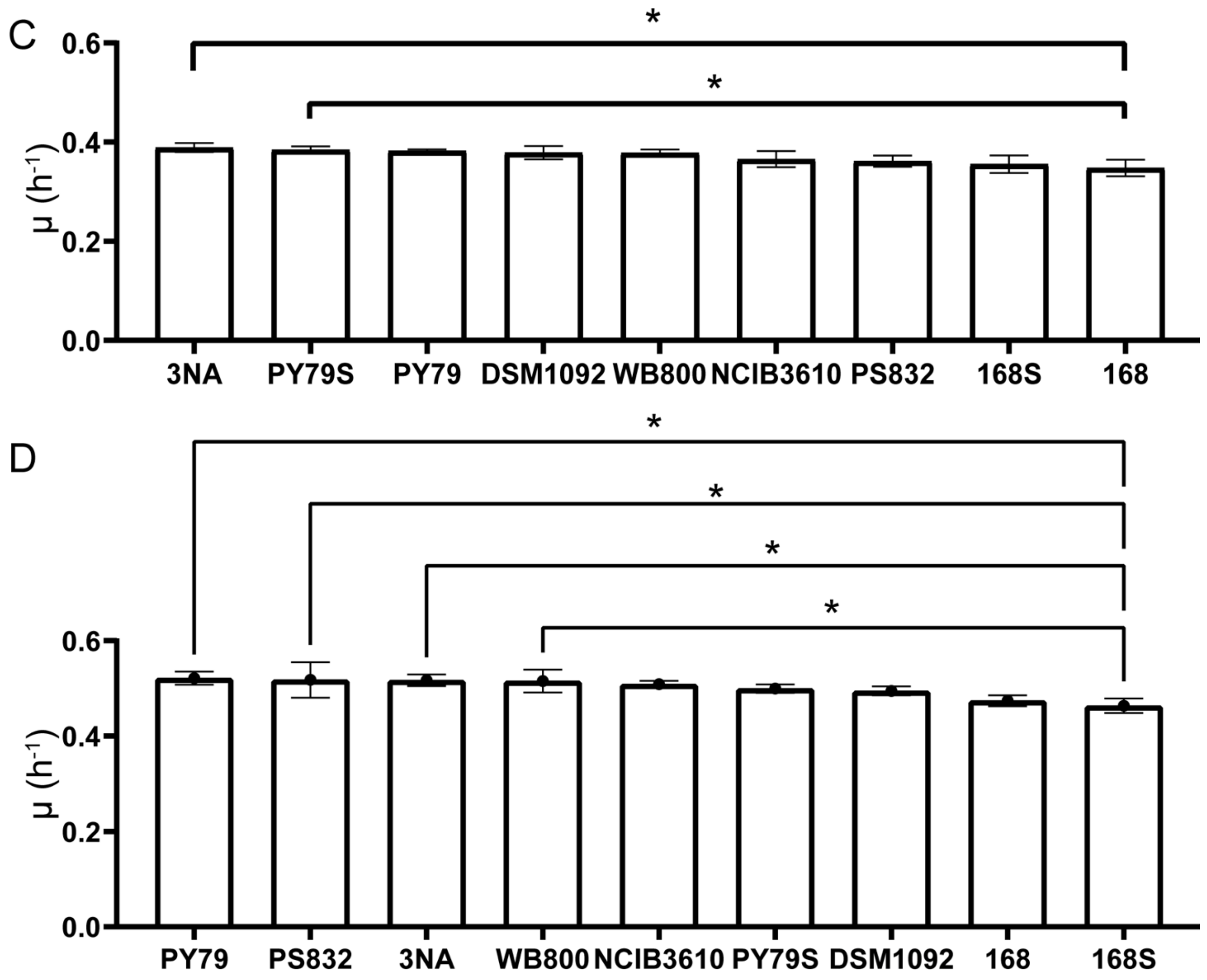


**Supplementary Figure 1 |** The pairwise one-way ANOVA analysis on the specific growth rates of nine strains on various media. A. Specific growth rate difference of nine strains in medium with ammonium sulphate; B**.** Specific growth rate difference of nine strains in medium with ammonium nitrate; C**.** Specific growth rate difference of nine strains with meat extract; D. Specific growth rate difference of nine strains in medium with meat extract and glucose. p <0.0001 (****), <0.001 (***), 0.01 (**) and <0.05 (*). ns: not significant.

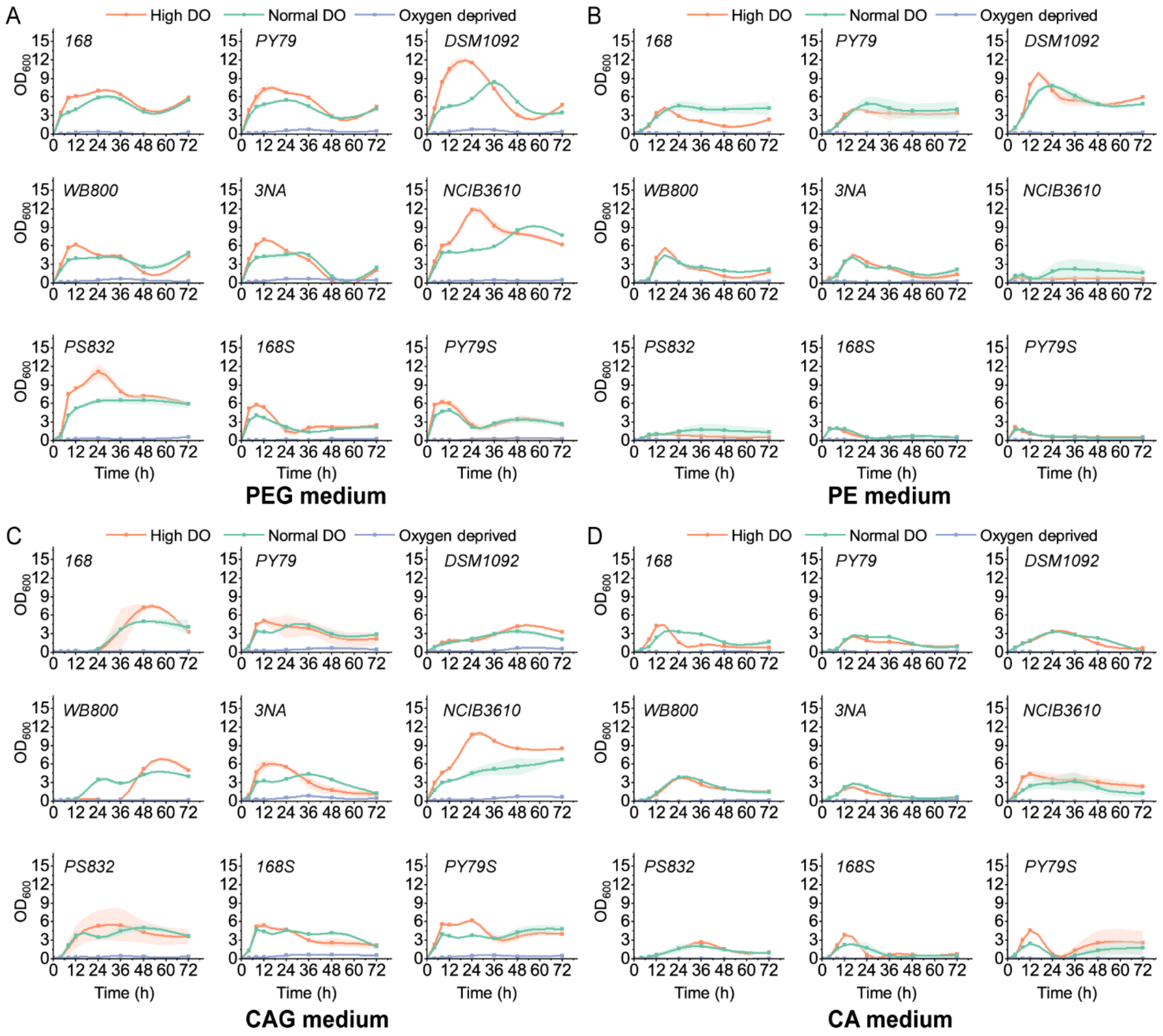


**Supplementary Figure 2 |** A~D: Growth curves of strains in flasks in medium with peptone and glucose (PEG), peptone (PE), casein hydrolysate and glucose (CAG) and casein hydrolysate (CA) under high DO, optimal DO and oxygen-deprived conditions. The shallow surrounding growth curve represents the standard deviations of replicates.

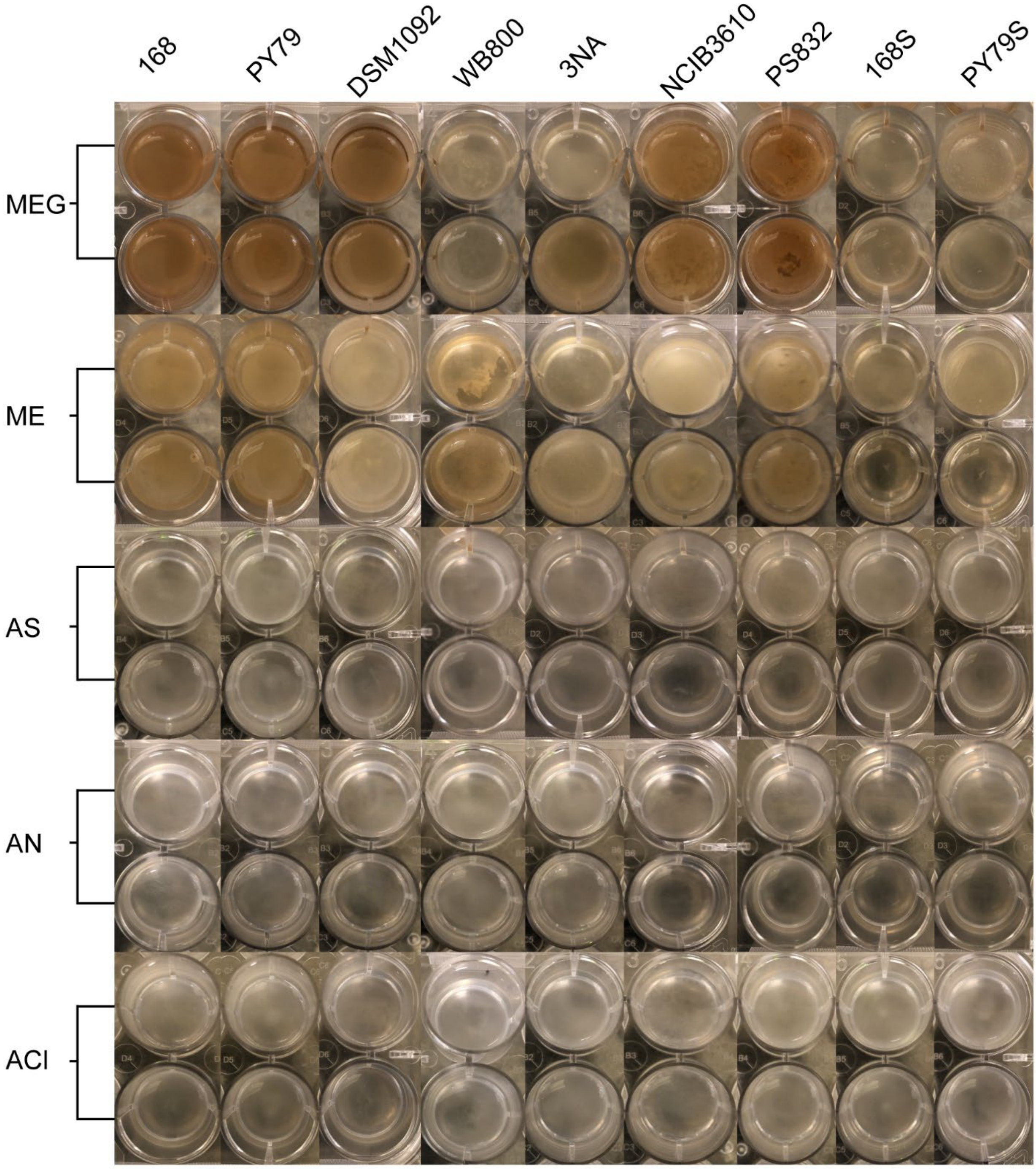


**Supplementary Figure 3 |** 1-day static incubation images of strains at 37°C in 24-well plates.

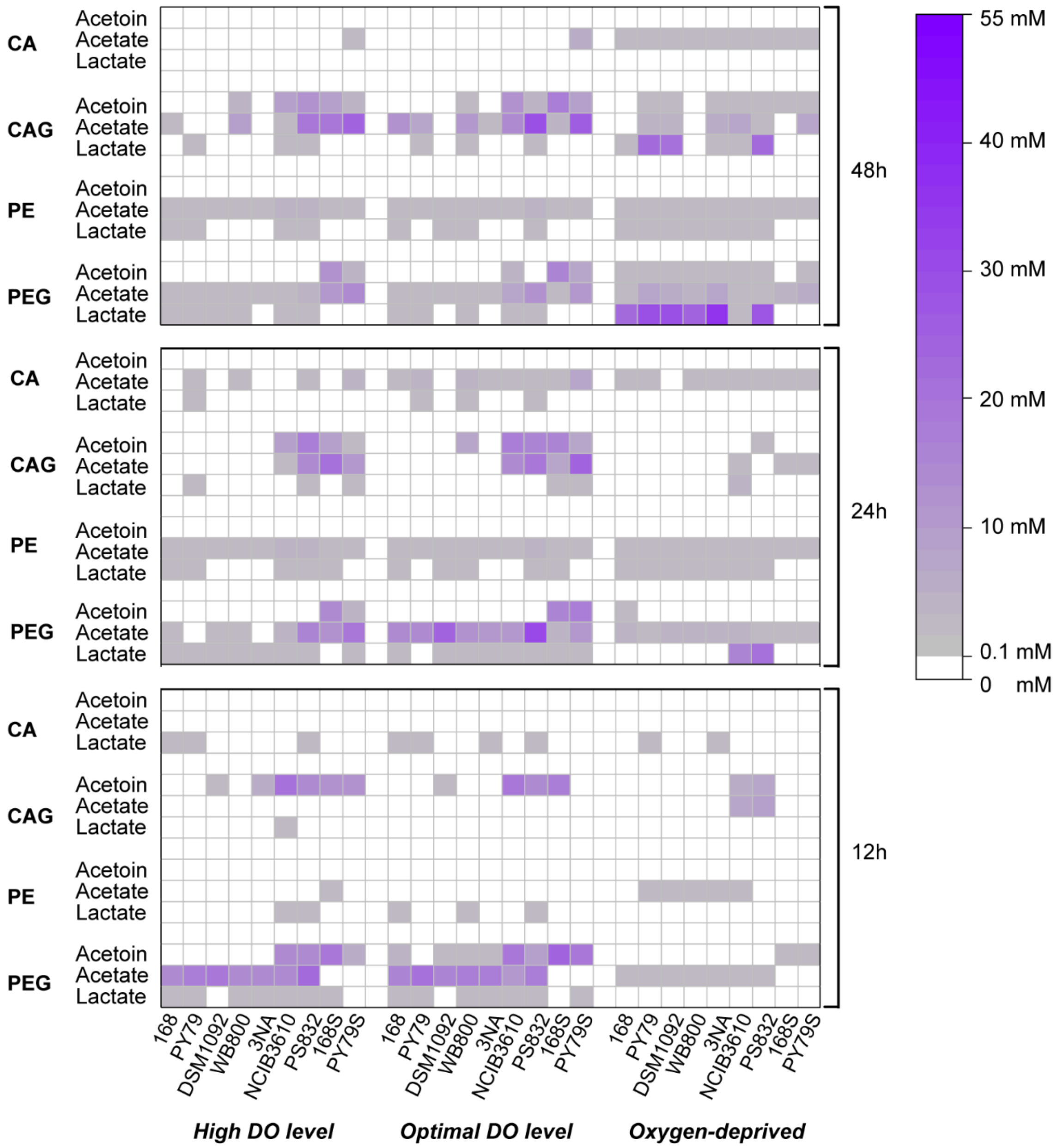


**Supplementary Figure 4 |** Metabolic distribution of nine *B. subtilis* strains in medium with casein hydrate (CA), peptone (PE), casein hydrate with glucose (CAG) and peptone with glucose (PEG) under high DO, optimal DO and oxygen-deprived conditions.
